# Physiological and behavioural characterisation of a novel steroid sulfatase-deficient mouse

**DOI:** 10.64898/2026.03.24.713857

**Authors:** Trevor Humby, Freya R. Shepherd, Talia Elgie, Libby Anderson-Watkins, Lucy I. Beevors, Angela E. Taylor, Paul A. Foster, William Davies

## Abstract

**Background:** Steroid sulfatase (STS) cleaves sulfate groups from steroid hormones. In humans, STS deficiency is associated with X-linked ichthyosis (a dermatological disorder), neurodevelopmental/mood conditions, and cardiac arrhythmias. Until recently, no single-gene ‘knockout’ mammalian model existed to investigate these associations; previous work in such a model has been limited to skin phenotypes.

**Methods:** We generated a novel C57BL/6J mouse model with a deletion in critical exon 2 of *Sts*. We then examined gene expression and enzyme activity in liver and brain samples of homozygous mice, and assessed the breeding performance and health of male and female deletion-carriers. Subsequently, we compared performance across a range of behavioural paradigms in wildtype and homozygous male and female mice: elevated plus maze, open field, rotarod, spontaneous alternation, and acoustic startle/prepulse inhibition. We also investigated serum steroid hormone levels by liquid chromatography-mass spectrometry and measured heart weights and two morphological indices (bodyweight/tibia length) *post mortem*.

**Results:** Homozygous mice almost completely lacked STS expression/activity. Genetically-altered mice exhibited grossly-normal breeding performance, health, and endocrinology. Homozygous mice were more active, and had higher normalised heart weights, than wildtype mice. We also found significant genotype x sex interactions on bodyweight, and on two behavioural measures (potentially reflecting lower anxiety in homozygous males and heightened anxiety in homozygous females).

**Conclusions:** The ‘*Sts*-deletion’ mouse represents an experimentally-tractable model in which to identify and characterise phenotypes associated with STS deficiency. The mechanistic basis of the genotype-phenotype associations described here requires further investigation, and whether such associations translate to humans remains to be tested.

## 1. Introduction

In mammals, the enzyme steroid sulfatase (STS) is solely responsible for cleaving sulfate groups from a variety of steroid hormones (e.g. dehydroepiandrosterone sulfate, DHEAS), thereby altering their water-solubility, activity, and availability as precursors in the biosynthesis of a number of oestrogens and androgens (Mueller et al., 2015). In humans, the *STS* gene is located at Xp22.31 and escapes X-inactivation; hence, expression and activity levels of the enzyme are higher in females than males and might influence sex-biased phenotypes (Davies, 2021). STS is widely-expressed, with highest levels in the placenta, arterial vasculature, adipose tissue, brain and gastrointestinal tract (GTex Portal, 2025; Mouse Genome Informatics, 2025a).

Loss of STS function in humans, typically occurring as a consequence of either a deletion Copy Number Variant or a disruptive gene-specific Single Nucleotide Variant (SNV), is often associated with X-linked ichthyosis (XLI), a rare dermatological disorder characterised by skin scaling as a consequence of cholesterol sulfate accumulation in the stratum corneum (Fernandes et al., 2010). Individuals affected by XLI, who are almost exclusively male, also appear at increased risk of numerous extracutaneous conditions, including cryptorchidism, corneal opacity, and fibrotic and haemostatic conditions (Wren and Davies, 2022). Female carriers of XLI-associated genetic variants may exhibit delayed or prolonged labour as a consequence of placental STS haploinsufficiency and impaired oestrogen-mediated cervical softening, but typically present with no, or mild, skin abnormalities (Fernandes et al., 2010).

Whilst gross cognitive function and academic attainment are typically unaffected in XLI, there is some evidence for mild-moderate impairment across aspects of fluid intelligence and memory (Brcic et al., 2020; Wren et al., 2024). Moreover, diagnostic rates of neurodevelopmental disorders (notably Attention Deficit Hyperactivity Disorder (ADHD), dyspraxia, autism and epilepsy) in XLI are 6-8 fold higher than in the general male population, and the fact that some individuals affected by neurodevelopmental conditions possess SNVs within *STS* provides support for loss-of-function of this gene as being causal (Wren and Davies, 2022). Rates of mood disorders and associated traits in males with XLI (and female carriers) also appear significantly elevated relative to their age and sex-matched counterparts from the general population (Wren and Davies, 2022). Genetic neuroimaging work has implicated smaller volume of the globus pallidus, a subregion of the basal ganglia, in vulnerability to neurodevelopmental and mood conditions in XLI (Wren et al., 2024). We have recently shown that individuals carrying genomic deletions encompassing *STS* are at substantially increased risk of cardiac arrhythmias than non-carriers (Brcic et al., 2020). Follow-up genetic association analyses have implicated STS deficiency as the most likely causal mechanism (Wren et al., 2023), and we have proposed that STS deficiency confers vulnerability to structural heart abnormalities (e.g. septal defects and ventricular hypertrophy) which in turn results in increased arrhythmia risk (Wren and Davies, 2024). Cardiac arrhythmias predispose to heart failure, stroke, accelerated cognitive decline/dementia, and potentially, in XLI, also to cardiac arrest (Davies, 2025). These conditions, like neurodevelopmental/mood conditions, can have long-term effects on health and morbidity/mortality, and burden healthcare systems. Characterising behavioural and cardiac phenotypes associated with STS deficiency, and identifying underlying pathophysiological mechanisms, will be valuable for enabling earlier identification of potential issues in individuals with XLI, and for developing targeted therapeutic strategies.

Traditionally, the physiological effects of loss of a single protein during mammalian development have been investigated through the use of knockout mice; the unique highly-repetitive genomic sequence within the mouse pseudoautosomal region has precluded such a strategy with respect to *Sts* and, as such, the impact of enzyme loss is poorly-defined. As an alternative to the knockout mouse, the 39,X^Y*^O mouse, which lacks *Sts* as a consequence of an end-to-end fusion of the X and Y chromosomes has been studied. 39,X^Y*^O mice recapitulate some features of neurodevelopmental conditions including inattention, altered response inhibition, aggression, hyperactivity, increased fluid consumption, heightened emotional reactivity and perseverative behaviours, and abnormal striatal and hippocampal serotonergic function (Davies et al., 2009; Trent et al., 2012a; Trent et al., 2012b; Davies et al., 2014). However, the 39,X^Y*^O model is necessarily male, is challenging to breed, and the deleted region includes multiple genes adjacent to *Sts* (including the autism candidate gene *Nlgn4*)(Trent et al., 2013; Kasahara et al., 2022); hence, the extent to which the aforementioned phenotypes are due to *Sts* deletion alone is unclear. The effects of selective loss of STS function in adulthood in rodents have also been investigated through acute administration of an enzyme inhibitor. These pharmacological studies have revealed effects on attention, behavioural inhibition, memory enhancement, hippocampal neurochemistry and neuroprotection in animal models relevant to dementia (Rhodes et al., 1997; Davies et al., 2009; Davies et al., 2014; Perez-Jimenez et al., 2021). Ideally, to understand the physiological role of STS, and the role of STS deficiency in the pathophysiology of XLI-related phenotypes, we require a mammalian model in which only STS is absent throughout development, and which can be easily-bred.

New technologies, such as CRISPR, through which a wide variety of genetic modifications can be readily made, have enabled the development of novel rodent models for understanding human disease (Bruter et al., 2024). Kwon and colleagues recently generated a mouse in which part of *Sts* exon 2 (encoding the enzyme’s sulfatase domain (Salido et al., 1996)) was deleted; their characterisation of skin morphology and biochemistry in this model implicated excessive generation of reactive oxygen species, dysregulation of Wnt/β and Hippo signalling pathways, and over-expression of E-cadherin as mechanisms contributing to XLI skin phenotypes (Kwon et al., 2024; Kwon et al., 2025). Here, we describe the generation, and initial physiological and behavioural characterisation, of an independent novel CRISPR-generated mouse model harbouring a small frameshift deletion in *Sts* exon 2, predicted to result in a premature stop codon. Guided by previous work in 39,X^Y*^O and STS-inhibition models, we initially focussed on basic sensory, motoric and exploratory paradigms. We also analysed serum steroid levels and normalised heart weights (a simple index of cardiac hypertrophy) in our new model. Given the early and exploratory nature of these investigations in this new model, a pre-registered protocol was not prepared.

## 2. Materials and Methods

### 2.1 Generation, genotyping, breeding and husbandry of mice

C57BL/6J-Sts^em2H^/H mice were obtained from the Mary Lyon Centre at MRC Harwell which is the UK node of the European Mouse Mutant Archive (EMMA) (www.infrafrontier.eu; repository number EM:15127). Mice were generated in collaboration with us as part of the Genome Editing Mice for Medicine (GEMM) programme according to the protocol in Mianne *et al*. (2017), and specific details are provided in **Supplementary Information**. Wildtype, heterozygous, and homozygous adult male and female mice were transported to Cardiff University, and initial breeding was conducted to generate heterozygous parents. Subsequently, heterozygous x heterozygous crosses were used to generate male and female wildtype, heterozygous and homozygous littermates. Mice were genotyped by Transnetyx (Cordova, TN) from genomic DNA obtained from ear biopsies at weaning using real-time PCR probes specific for wild type and mutant alleles; eight mice were re-genotyped *post mortem* and results were identical to previous calls. Mice were housed in same-sex mixed-genotype groups (2-5 animals per cage), and were maintained on *ad libitum* food and water, in a temperature, humidity and light-controlled room (21±2 °C, 50±10% humidity, lights on at 0700hr for 12h) with environmental enrichment, and were regularly inspected for signs of ill health. Animals of each genotype for the various analyses came from ≥6 homecages and individual animals were the experimental unit. Unless otherwise stated, researchers were aware of animals’ genotypes prior to testing; most measures were obtained automatically, or were blindly double-scored, to minimise potential researcher bias. Animals of the various genotypes were tested in a randomised order based upon their number assigned at weaning (i.e. prior to genotypes being known). Experiments were performed according to the UK Animal Scientific Procedures Act (1986) under Home Office Project Licences P825DDA5B, PP2050169 and PP6583178 and comply with the ARRIVE guidelines.

### 2.2 Behavioural analysis

92 adult (3-5months) mice (wildtype (+/+) males (n=12), heterozygous (+/-) males (n=15), homozygous (-/-) males (n=22), wildtype females (n=14), heterozygous females (n=15), and homozygous females (n=14)) underwent behavioural testing conducted by a female researcher during the light phase (0900-1500hrs). For the sake of clarity the heterozygote data are not reported here, but across the vast majority of measures, these groups presented, as expected, intermediately between wildtype and homozygote groups. This sample size allowed medium-large effect sizes (f>0.36) between wildtype and homozygous animals to be reliably detected (with >80% power, α=0.05); such effect sizes are comparable with those we have seen previously when comparing 40,XY and 39,X^Y*^O mice (Trent et al., 2012a).

Following two weeks’ habituation to handling, a series of behavioural tests assaying anxiety-related behaviours, locomotor activity, motoric function/co-ordination, exploratory behaviours and startle response/sensorimotor gating was conducted across three days, with the least stressful tests taking place first and with at least 90 minutes between tests to allow stress responses to return to baseline. On each test day, male mice were run before female mice, with animals run in a pseudorandomised order with respect to genotype. Test apparatus was cleaned with 70% industrial methylated spirits between animals to disinfect it and to limit any potential confounding effects of odour. Mice were allowed to habituate to each testing room for ≥30mins prior to testing. The order of testing was: elevated plus maze (EPM), open field test (OFT), assessment of consummatory behaviour, rotarod, spontaneous alternation and finally the assessment of acoustic startle response and sensorimotor gating.

The EPM and OFT were run under identical lighting conditions (15 lux) and the position of each mouse in the apparatus was automatically tracked using EthoVision XT software (Noldus Information Technology, Netherlands) at a rate of 12 frames/s via a camera mounted above the centre of each piece of apparatus. The EPM was constructed of white Perspex and consisted of four arms of equal size (175×78mm) extending from a central square region (78×78mm) and positioned 450mm above the floor. Two of the arms were ‘open’ (no walls) and two were enclosed by 150mm high opaque walls (‘closed’). Arms of the same type were diametrically opposed. Animals were placed in the same enclosed arm, facing the wall, at the start of the trial, and were allowed to freely explore the apparatus for five minutes. Key EPM outcome measures assaying anxiety-related behaviours and/or within-maze activity included: the ratio of time spent in the open and closed arms (x100), total distance moved, latency to enter the open arm and number of fecal boli deposited. The OFT comprised a 750×750mm white Perspex arena with 450mm high walls. Animals were placed in the same corner of the arena, facing the wall, and were allowed to explore freely for 10 minutes. The arena was divided into two concentric virtual zones: the ‘inner zone’ (central 600×600mm square) and the ‘outer zone’. Key OFT outcome measures assaying anxiety-related behaviours and/or within-maze activity included: time in the inner zone, total distance moved, latency to enter the inner zone, and number of fecal boli deposited.

To assess baseline drinking behaviour and response to a novel foodstuff, mice with previous *ad libitum* access to homecage water were individually placed into a small chamber (285×130×120mm, length x width x height) for 10mins in which two small containers (maximum volume ~3ml each) were positioned towards the rear, one containing tap water and the other a 10% condensed milk solution (Nestle Ltd) novel to the mouse. Containers were weighed at the start and end of the session and the total fluid volume consumed by each animal calculated; this figure was then normalised to bodyweight^0.75^ to take into account Kleiber’s 0.75 mass exponent and multiplied by 100. The volume of milk consumed as a percentage of total fluid consumed was also calculated.

Motor learning and co-ordination were assessed using a mouse rotarod (Model 47600, Ugo Basile, Italy). The rod (30mm diameter) was coated with rubber grooves to provide grip. Each animal completed five trials of 300s maximum duration with ≥30mins recovery time between each trial. During each trial, the rotation speed increased from 5–50 rpm at a constant rate of 0.15rpm/s. The latency to fall was recorded on each trial, when the mouse caused a timer to stop when triggered by a lever (160mm below the rod). For each animal, the median time spent on the apparatus before falling across the five trials was determined.

Exploratory and spontaneous alternation behaviour, partially indexing working memory ability, was assayed by testing animals in a T-maze made of clear Perspex with a ‘start arm’ 54.5cm long and 10cm wide, two ‘goal arms’ 25.5cm long and 10cm wide, a ‘decision’ space 10cm × 10cm, and walls 25cm high. Mice were initially placed on the floor at the end of the ‘start’ arm and were allowed to explore freely for 10mins with their behaviour being video-recorded. To enhance between-arm discriminability, the right arm was covered in a checkerboard pattern (alternating squares 2.5 × 2.5cm) and the left arm covered in a dot pattern (black dots of 1cm diameter). Key outcome measures included: total arm entries (a measure of locomotor activity defined by all four paws in an arm) and total alternations (defined by consecutive visits to the three different arms). These measures were initially scored from videos by an experienced analyst blind to animals’ genotype and sex, and were double-scored by a second blinded researcher. The ratio of spontaneous alternations to maximum possible spontaneous alternations (i.e. total arm entries-2) was then calculated. Inter-rater reliability for the key measures was determined via calculation of the intra-class correlation coefficient (ICC).

Acoustic startle response (ASR) and prepulse inhibition (PPI) of this, indices of reactivity to an unpredictable auditory stimulus and sensorimotor gating respectively, were measured using apparatus from SR-Lab (San Diego Instruments, USA). Animals were placed in a clear Perspex tube (35mm internal diameter) mounted on a Perspex plinth in a sound-attenuating chamber. A 70dB background white noise stimulus was continuously played throughout the session via a loudspeaker positioned 120mm above the tube. The whole-body startle response was detected on each trial by a piezoelectric sensor attached to the plinth, which transduced flexion in the plinth into a digitised signal. The session comprised 44 trials in total, following a 5 minute habituation period with background sound and lasted for approximately 30 minutes. The startle response was recorded in arbitrary startle units using SR-Lab software over the 65ms period following the main startle stimulus onset. Acoustic stimuli were presented with a mean intertrial interval of 16s (pseudorandomly varied between 6 and 24 seconds). Each pulse-alone trial consisted of a 40ms 120dB startle stimulus. The median startle response for each mouse across the final 8 pulse-alone trials was calculated and the resultant figure divided by bodyweight and multiplied by 10. Prepulse trials consisted of a 20ms prepulse stimulus (4, 8, or 16dB) followed by a 40ms 120dB startle stimulus occurring 80ms after prepulse offset. There were six trials at each prepulse amplitude randomly interspersed amongst the pulse-alone trials. PPI was calculated as the percentage reduction in mean startle amplitude between prepulse and pulse-alone trials.

### 2.3 Tissue collection

80 behaviourally-tested, and behaviourally-naïve, adult (3-11.5 months) mice were weighed and culled by cervical dislocation between 09:00-15:00hrs, with the culling order pseudorandomised by genotype (wildtype males (n=15), homozygous males (n=29), wildtype females (n=21), homozygous females (n=15)). Trunk blood was collected and serum extracted using BD Microtainer SST tubes (Fisher Scientific) for steroid analysis. Whole brain and liver samples were also taken for enzyme activity and gene expression analyses (n=2 per group). Serum, brain and liver samples were snap-frozen on dry ice and stored at −80°C. Hearts from older animals (>20 weeks) were dissected, residual blood removed, and their wet weight recorded; tibias from these animals were also dissected, and their length measured within 0.1mm using digital vernier calipers. For each animal, heart weight was divided by bodyweight and tibia length, and the resultant figure multiplied by 1000. For the bodyweight and heart weight measures, the study was powered to reliably detect (>80% power) a genotype-dependent medium effect size (f>0.35, α=0.05).

### 2.4 Enzyme activity and gene expression analyses

STS activity in liver and whole brain tissues was assayed by following our previously-published cell-free method with minor modifications for solid samples (Gilligan et al., 2017); details are provided in **Supplementary Information**. Individual animals’ samples were assayed in triplicate and the median value taken for statistical analysis. Additionally, *Sts* gene expression in liver samples from homozygous and wildtype mice was assayed using quantitative PCR (methods in **Supplementary Information**).

### 2.5 Serum steroid analysis

Serum steroid levels were quantified at the Steroid Metabolome Analysis Core, University of Birmingham, using a validated assay described previously (Schiffer et al., 2022); details are provided in **Supplementary Information**.

### 2.6 Statistics

Statistical analysis was performed with IBM SPSS Statistics 27 software. Categorical breeding data were analysed by chi-squared test. Following removal of group outliers (<1.5 times the interquartile range below the 25^th^ percentile or above the 75^th^ percentile, <3.5% of all datapoints), most behavioural and cardiac morphology data were analysed by Two Way ANCOVA with factors of GENOTYPE (wildtype or homozygote) and SEX (male or female), and age as a covariate; estimated marginal means are reported for significant interaction effects. Analysis of the prepulse inhibition data also included the repeated measures factor of PREPULSE AMPLITUDE (4, 8 or 16dB) with Greenhouse-Geisser correction to adjust for lack of sphericity. Where assumptions underlying ANCOVA were violated (non-normal distribution of residuals in ≥two groups assessed by Shapiro-Wilk test, or non-homogeneity of variance assessed by Levene’s test), data were appropriately transformed as far as possible (natural log or reciprocal). For analyses where ANCOVA assumptions were significantly violated (e.g. for steroid hormone analysis), comparisons were conducted by Mann-Whitney U-test or Kruskal-Wallis H-test with follow-up Bonferroni-corrected pairwise comparisons to adjust for multiple testing. P-values<0.05 were regarded as being statistically-significant. Data are shown as box and whisker plots where centre lines indicate median values, box limits indicate the 25th and 75th percentiles and whiskers extend 1.5 times the interquartile range from the 25th and 75th percentiles; outlying datapoints reside outside the plot whiskers.

## 3. Results

### 3.1 Effect of genetic manipulation on STS enzyme activity and gene expression

Consistent with previous mouse data (Nicolas et al., 2001), STS activity was higher in wildtype liver tissue (313.3±105.3pmol/mg/hr, n=2 per sex) than in wildtype brain tissue (53.2±9.4pmol/mg/hr, n=2 per sex). As expected, the genetic deletion resulted in substantial attenuation of STS activity in homozygous mice across both tissues. In liver, STS activity was barely detectable in homozygous animals (n=2 per sex) at 1.83±1.83pmol/mg/hr (>99% reduction compared to wildtype animals, U=16.0, p=0.03). In brain, STS activity in homozygote animals was just 5.6±4.1pmol/mg/hr (>85% reduction compared to wildtype animals, U=16.0, p=0.03). These enzyme activity data mirrored our gene expression data, where *Sts* gene expression in homozygous mouse liver tissue was markedly lower (~15-fold) than that in wildtype tissue (**Supplementary Figure 1**).

### 3.2 General health, bodyweight and breeding performance

The vast majority (>95%) of male and female heterozygous and homozygous mice exhibited normal health from birth to the latest timepoint assessed (12 months). Of the 300 adult genetically-altered mice (heterozygotes or homozygotes) generated in our work to date, three weaned heterozygous mice (two male, one female) were found dead of an indeterminate cause (1.0%) and one female homozygous mouse required culling due to severe illness of uncertain origin (0.3%). Two heterozygous, and one homozygous, adult female mice were identified as having vaginal septa (2.1% of genetically-altered females), and of the 65 genetically-altered female mice used for breeding to date, four heterozygotes (6.2%) exhibited evidence of dystocia. Measurement of bodyweight across animals aged 98-348 days revealed no significant age-adjusted main effect of GENOTYPE (F[1,85]=0.23, p=0.63); however, there was the expected significant effect of SEX (males>females)(F[1,85]=259.7, p<0.001) and an unexpected significant GENOTYPE x SEX interaction, whereby homozygous males were ~5% lighter (estimated marginal mean=32.5±0.4g), and homozygous females ~5% heavier (estimated marginal mean=25.8±0.6g), than their sex-matched wildtype littermates (estimated marginal means of 34.0±0.5g and 24.7±0.5g respectively) (F[1,85]=6.20, p=0.02)(**Figure 1**).

**Figure 1.**
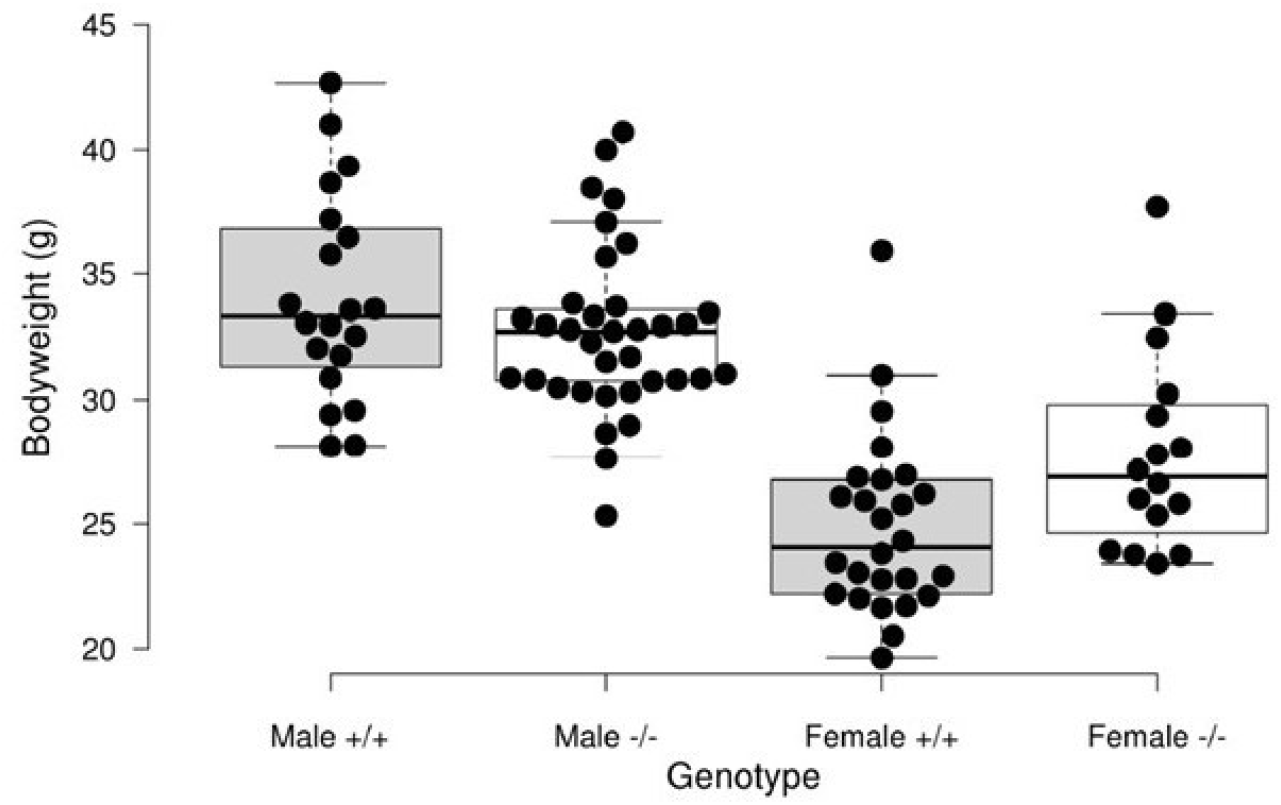
Bodyweight at culling in adult male and female wildtype (+/+) and homozygous (-/-) mice

To date in our breeding colony, 303 mice from 54 litters have survived to weaning from heterozygote x heterozygote crosses (median litter size of six pups). At weaning, the genotype ratios obtained were consistent with the expected Mendelian ratios (wildtype male (n=31, 10%), heterozygote male (n=93, 31%), homozygote male (n=45, 15%), wildtype female (n=33, 11%), heterozygote female (n=73, 24%) and homozygote female (n=28, 9%)(χ^2^[5]=9.8, p=0.08). The genotypes of 26 unsexed pups found dead between birth and weaning were also in the expected ratios (wildtype (n=4, 15%), heterozygote (n=14, 54%), homozygote (n=8, 31%)(χ^2^[2]=1.38, p=0.50), suggesting no large effect of genotype on deaths in this period.

### 3.3 Behavioural analysis

#### 3.3.1 Elevated plus maze test

On the main measure of anxiety-related behaviour in this task (ratio of time spent on the open arms relative to time spent in the enclosed arms), there was no main effect of GENOTYPE (F[1,55]=0.44, p=0.51), but there was a significant main effect of SEX, whereby females tended to spend more time on the open arms relative to males (F[1,55]=4.70, p=0.03)(**Figure 2A**). There was also a GENOTYPE x SEX effect on this measure, with homozygous males tending to spend more time on the open arms (estimated marginal mean=132.6±13.7), and homozygous females tending to spend less time on the open arms (estimated marginal mean=115.5±18.6), than their wildtype sex-matched controls (estimated marginal means of 67.2±18.5 and 158.1±17.2 respectively)(F[1,55]=10.0, p=0.003)(**Figure 2A**). On the ‘latency to enter the open arm’ measure (**Figure 2B**), there was a significant main effect of GENOTYPE, with homozygous animals tending to enter the open arms sooner in the test than wildtype mice (F[1,56]=4.23, p=0.04); the main effect of SEX on this measure was non-significant (F[1,56]=3.32, p=0.07) as was the GENOTYPE x SEX interaction (F[1,56]=3.25, p=0.08). There were no significant group effects on a final measure of anxiety-related behaviour in this test: number of fecal boli deposited (H[3]=0.35, p=0.95). The significant effects on anxiety-related behaviour described above occurred in the absence of significant effects of GENOTYPE (F[1,55]=0.13, p=0.72), SEX (F[1,55]=0.74, p=0.39), or GENOTYPE x SEX (F[1,55]=0.58, p=0.45) on activity (total distance moved within the maze) (**Figure 2C**). Covarying for ‘distance moved’ did not impact the pattern of results with respect to open:closed arm time, but resulted in attenuation of the genotype-dependent effect on ‘open arm entry latency’ (F[1,55]=3.95, p=0.052) implying its partial dependence upon general activity levels.

**Figure 2.**
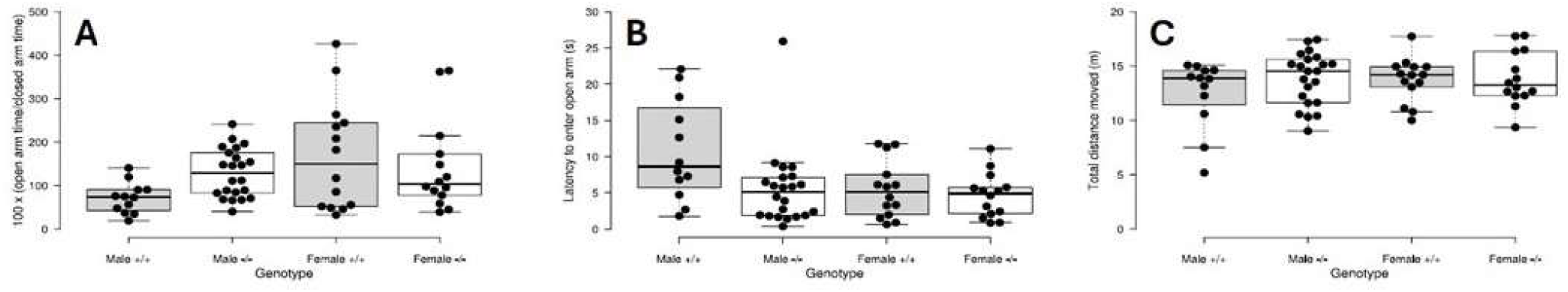
Performance of adult male and female wildtype (+/+) and homozygous (-/-) mice on key measures of anxiety-related behaviour (open:closed arm time (A) and latency to enter open arm (B)) and activity (total distance travelled) on the elevated plus maze test.

#### 3.3.2 Open field test

We observed no significant effects of GENOTYPE (F[1,57]=1.14, p=0.29), SEX (F[1,57]=2.77, p=0.10) or GENOTYPE x SEX (F[1,57]=1.74, p=0.19) on the main measure of anxiety-related behaviour (time spent in the central zone)(**Figure 3A**). A similar null pattern of findings was observed with respect to a second measure of anxiety-related behaviour (latency to first enter the central zone): effect of GENOTYPE (F[1,55]=0.19, p=0.67), effect of SEX (F[1,55]=0.10, p=0.75) and GENOTYPE x SEX (F[1,55]=0.33, p=0.57)(**Figure 3B**). There were also no significant group effects on a third measure of anxiety-related behaviour in this test: number of fecal boli (H[3]=1.44, p=0.70). We did, however, identify a genotype-dependent effect on locomotor activity (total distance moved) in this relatively non-aversive, unrestricted test (F[1,56]=4.19, p=0.045) with homozygotes being ~10% more active than their wildtype counterparts. There was no significant main effect of SEX (F[1,56]=0.004, p=0.95), nor a significant GENOTYPE x SEX interaction (F[1,56]=0.13, p=0.72) with respect to open field activity (**Figure 3C**).

**Figure 3.**
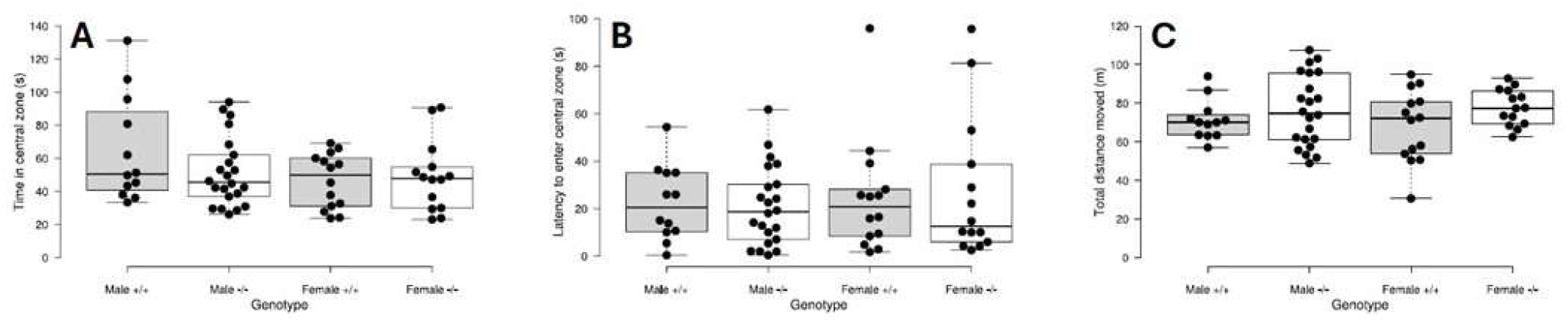
Performance of adult male and female wildtype (+/+) and homozygous (-/-) mice on key measures of anxiety-related behaviour (time in central zone (A) and latency to enter central zone (B)) and activity (total distance travelled (C)) on the open field test.

#### 3.3.3 Baseline drinking behaviour and reactivity to a novel foodstuff

There was a significant overall group effect on total fluid volume consumed normalised to bodyweight^0.75^ (H[3]=8.51, p=0.04). This effect appeared due to a combination of higher consumption in homozygous male mice relative to wildtype male mice (resembling the effect reported in 39,X^Y*^O mice previously, Trent et al., 2012a) and slightly lower consumption in homozygous female mice relative to wildtype female mice; however, no individual adjusted pairwise comparisons were significant (**Supplementary Figure 2A**). With respect to solution preference (water vs. milk), there was no significant effect of GENOTYPE (F[1,57]=0.20, p=0.65) or SEX (F[1,57]=2.44, p=0.12), but there was a significant GENOTYPE x SEX interaction (F[1,57]=5.69, p=0.02) mirroring the baseline drinking pattern, with homozygous males exhibiting a greater preference for the milk solution (estimated marginal mean=61.4±4.4) than sex-matched wildtype mice (estimated marginal mean=50.9±6.0), and homozygous females exhibiting a reduced preference for the milk solution (estimated marginal mean=40.0±5.6) relative to sex-matched wildtype mice (estimated marginal mean=55.4±5.6)(**Supplementary Figure 2B**). Across the four groups, there was no significant group effect on number of fecal boli (H[3]=3.13, p=0.37).

#### 3.3.4 Rotarod test

Median time on the rotarod across the five trials was equivalent across the two genotypes (F[1,57]=0.94, p=0.34) and the two sexes (F[1,57]=0.33, p=0.57), and there was no significant GENOTYPE x SEX interaction (F[1,57]=3.49, p=0.07)(**Supplementary Figure 3**). The number of fecal boli deposited on the final trial did not differ by group (H[3]=2.15, p=0.54).

#### 3.3.5 Spontaneous alternation test

The behavioural performance of a subset of 30 animals split across genotypes and sexes was independently, and blindly, double-scored for key measures. There was very good-excellent, and highly-significant (p<0.001), agreement between blinded raters: intra-class correlation coefficients=0.97 for ‘total arm entries’, 0.87 for ‘spontaneous alternations’, and 0.81 for ‘spontaneous alternations as a function of maximum possible alternations’.

There was a highly-significant effect of GENOTYPE on the total number of spontaneous alternations (F[1,55]=15.29, p<0.001) reflecting higher numbers of alternations in homozygous mice, but no effect of SEX (F[1,55]=1.94, p=0.17) and no GENOTYPE x SEX interaction (F[1,55]=1.13, p=0.29)(**Figure 4A**). Similarly, there was a significant effect of GENOTYPE on total arm entries (F[1,55]=4.56, p=0.04) consistent with more arm entries in homozygous mice, but there was no significant effect of SEX (F[1,55]=0.14, p=0.71) or GENOTYPE x SEX (F[1,55]=2.62, p=0.11) observed (**Figure 4B**). There was no significant overall group effect on the proportion of alternations as a function of the number of maximum possible alternations (H[3]=6.11, p=0.11)(**Figure 4C**). There was also no overall group effect on the number of fecal boli deposited (H[3]=5.05, p=0.17).

**Figure 4.**
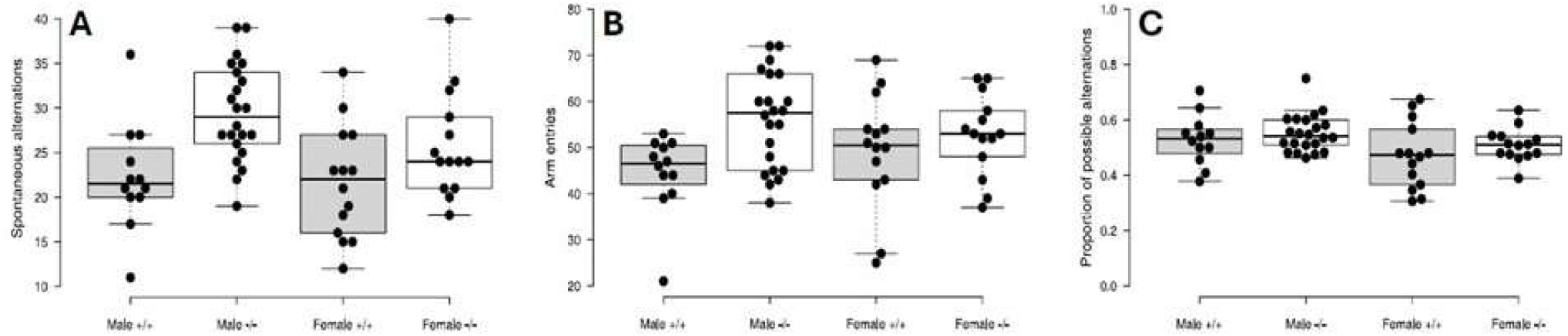
Performance of adult male and female wildtype (+/+) and homozygous (-/-) mice on alternation (A), activity (B) and alternations normalised for activity (C) on the spontaneous alternation test

#### 3.3.6 Acoustic startle response and prepulse inhibition

There was no significant effect of GENOTYPE (F[1,52]=1.22, p=0.28), SEX (F[1,52]=1.15, p=0.29) or GENOTYPE x SEX (F[1,52]=1.69, p=0.20) on the basic startle response normalised for bodyweight (**Supplementary Figure 4A**). Although the expected increase in startle inhibition was seen with increasing PREPULSE AMPLITUDE across groups (F[1.51,87.7]=93.3, p<0.001), there was no significant effect of GENOTYPE (F[1,57]=2.06, p=0.16), SEX (F[1,57]=0.53, p=0.47), or GENOTYPE x SEX (F[1,57]=0.28, p=0.60) on prepulse inhibition (**Supplementary Figure 4B**).

### 3.4 Steroid hormone analysis

21 steroids in total were assayed in serum across four groups: wildtype males (n=6), wildtype females (n=5), homozygous males (n=15) and homozygous females (n=3). The four groups did not differ significantly by age (H[3]=4.21, p=0.24). Eight steroids (pregnenolone, 17α-hydroxyprogesterone, 11-deoxycorticosterone, androstenedione, 11-ketoandrostenedione, 11-ketotestosterone, 11β-hydroxyandrostenedione, 11β-hydroxytestosterone) were not detectable in >10% of all samples, with lack of detectability not differing significantly by group for seven of these (0.13<p<1.0); androstenedione was less frequently detectable in female than male samples (~33% vs. ~80%, p=0.04). Levels of the remaining 13 steroids were undetectable in <1.5% of all sample readings, where they were assigned a zero value. For testosterone, dihydroepiandrosterone and 5α-androstanedione, there were significant overall and pairwise effects across the four groups, consistent with the expected higher levels in male compared to female groups, but not indicative of genotype-dependent effects (**Table 1**). For aldosterone, there was a significant effect of group, but no significant pairwise comparisons. For the other readily-detectable compounds assayed, there was no evidence for either genotype or sex-dependent effects.

**Table 1.**
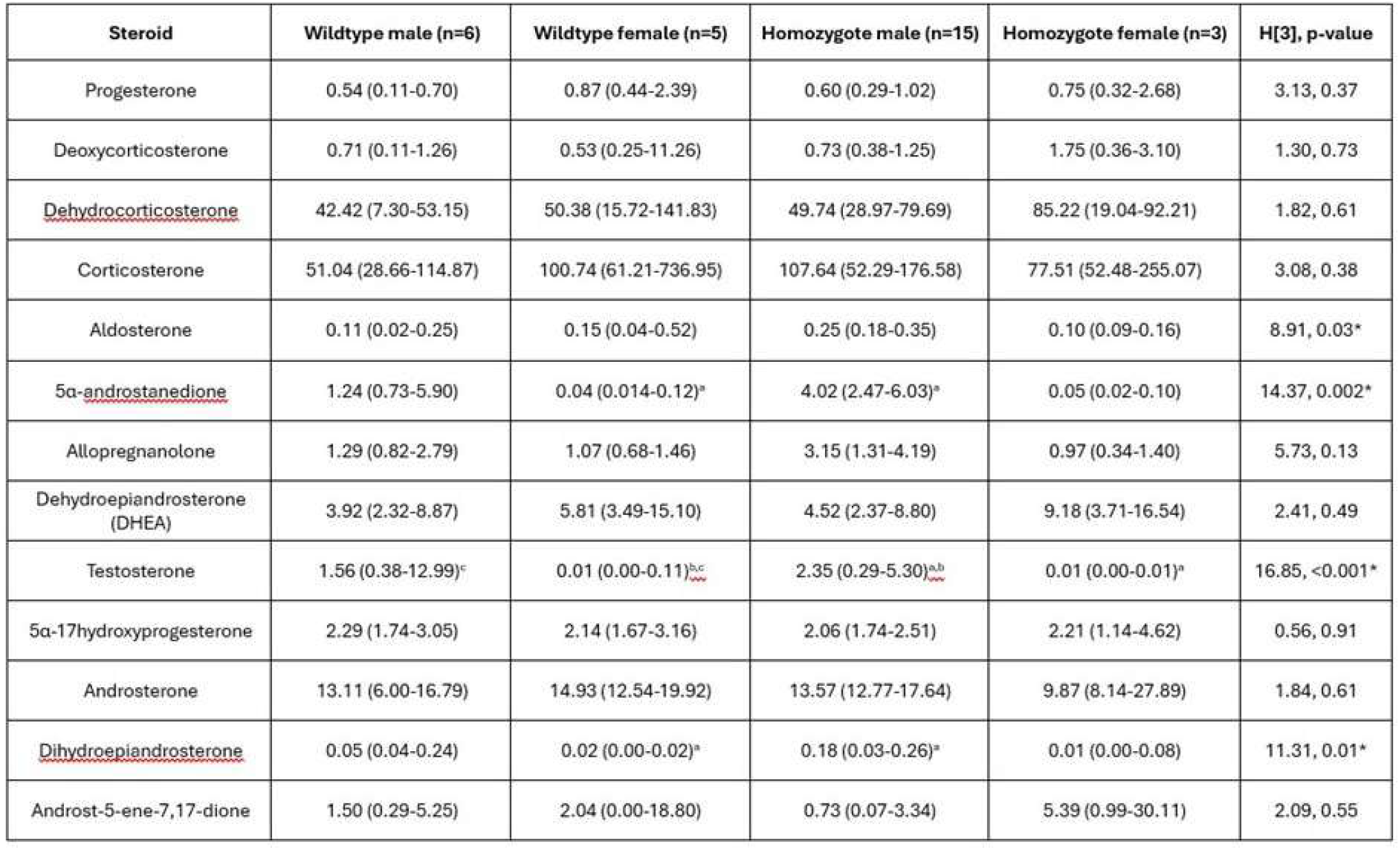
Median concentrations (ng/ml) of serum steroid hormones across groups with 95% confidence intervals defined by bootstrapping. Asterisks indicate significant (p<0.05) overall group effects, and subscript letters a-c denote significant (p<0.05) pairwise comparisons.

### 3.5 Normalised heart weights

In older mice (>20 weeks), where heart pathology would be expected to be greatest, there was a significant age-adjusted effect of GENOTYPE on heart weight:bodyweight (F[1,67]=4.91, p=0.03), but there was no significant effect of SEX (F[1,67]=2.72, p=0.10) nor any significant GENOTYPE x SEX interaction (F[1,67]=2.23, p=0.14)(**Figure 5A**). With respect to the heart weight:tibia length measure (**Figure 5B**), after covarying for age, there was a highly-significant effect of GENOTYPE (F[1,62]=10.78, p=0.002, reflecting a ~10-20% greater ratio in homozygous animals compared to wildtype animals), as well as a highly-significant effect of SEX (males>females)(F[1,62]=109.15, p<0.001), consistent with previous data on this measure in C57BL/6J mice (Gurgen et al., 2011). There was no significant GENOTYPE x SEX interaction on the heart weight:tibia length ratio (F[1,62]=0.13, p=0.72).

**Figure 5.**
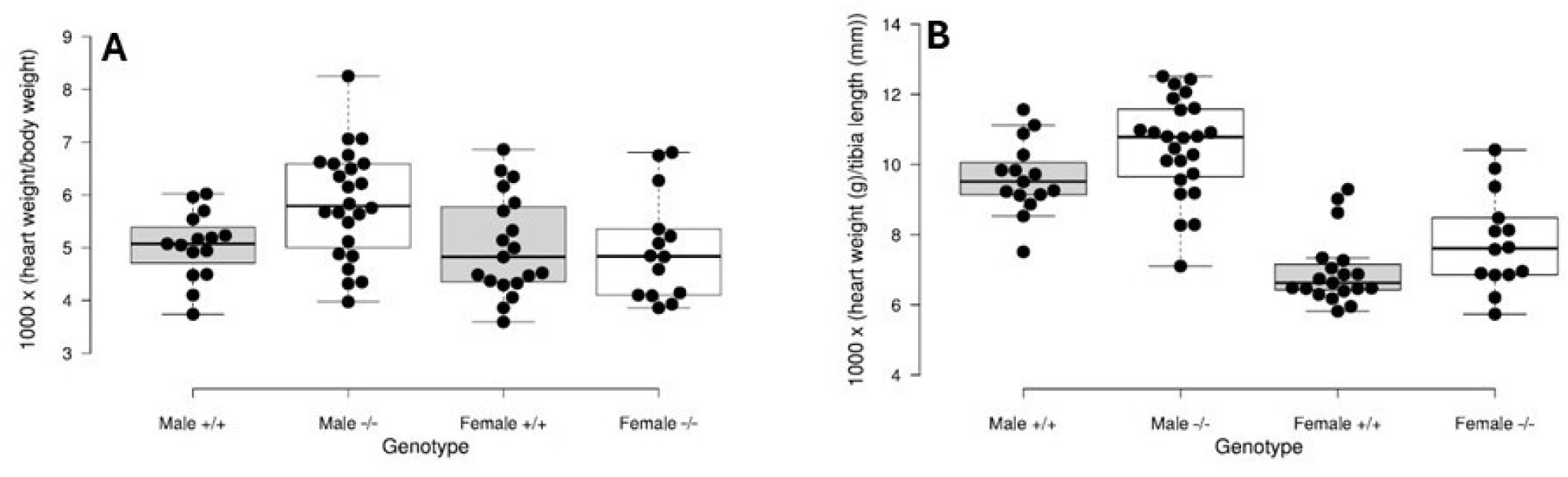
Heart weight relative to two morphological features (bodyweight (A) and tibia length (B)) in older (>20 weeks) adult male and female wildtype (+/+) and homozygous (-/-) mice

## 4. Discussion

The steroid sulfatase enzyme, encoded by the sex-linked *STS* gene, catalyses the conversion of sulfated to free steroids. In humans, STS deficiency is associated with a range of physiological and behavioural phenotypes, including X-linked ichthyosis. Although skin phenotypes have been investigated previously in mice with a null mutation within *Sts*, the consequences of STS deficiency on general health, physiology and behaviour in mice have not yet been described.

We generated a novel genetic mouse model in which significantly attenuated STS expression and activity were predicted. We confirmed that this was the case in central (brain) and peripheral (liver) tissues of homozygous mice, although there was some degree of residual STS activity in the former; the efficiency of murine nonsense-mediated decay processes can differ by tissue (Zetoune et al., 2008).

Heterozygous females gave birth to litters of equivalent size to those seen in wildtype mice of the C57BL/6 background strain (Mouse Genome Informatics, 2025b) and in the expected genotype ratios, and mice lacking STS activity exhibited grossly normal health until at least 12 months of age. These data suggest that STS deficiency in mice, as in humans, has no major effects on general health or mortality. Reproductive health issues (vaginal septa and dystocia) were noted in genetically-altered female mice. The prevalence of the former issue was comparable to that seen in wildtype mice of the same strain (<5%) (The Jackson Laboratory, 2025), and, as such does not appear to be genotype-dependent. The prevalence of dystocia in (functionally) wildtype C57BL/6 mouse mothers is ~0-8% (Kalev-Altman et al., 2023; Levenson et al., 2020; Marx et al., 2013), and in wildtype members of other multiparous small mammalian species is ~5-8% (Kutzler, 2008); the prevalence we observed in heterozygous mice (~6%) is within this normal range. However, given that heterozygosity for *STS*-null variants is associated with delayed or prolonged labour in humans, this phenotype should remain an area of focus as more data are collected from further breeding pairs. To minimise any potential welfare issues, younger heterozygous mice (<5 months of age) should be used for breeding.

Our preliminary finding of lower bodyweight in STS-deficient males compared to sex-matched wildtype controls, but higher bodyweight in STS-deficient females compared to sex-matched wildtype controls is interesting and worthy of follow-up. Should it be replicated in additional experimental cohorts, its mechanistic basis might be investigated further e.g. examining whether the differences arise due to genotype-dependent effects on muscle mass, levels of adiposity, food intake, metabolism and/or activity levels. To date, no systematic studies have been conducted examining bodyweight in humans lacking STS; if our findings are robust and translatable, we might expect males diagnosed with XLI to be lighter than their unaffected peers, and female carriers to be heavier than equivalent non-carriers, and for the mechanistic basis of any phenotypes to be conserved across species.

Behaviourally, the most consistent positive finding was a small, but significant, genotype-dependent effect on general activity measures across two tests (distance travelled in the open field, and number of arm entries in the spontaneous alternation paradigm); elevated activity levels as a consequence of STS deficiency may contribute to the hyperactivity phenotype described previously in 39,X^Y*^O mice (Trent et al., 2012a). This finding supports the idea that STS deficiency in humans predisposes to increased hyperactive-impulsive traits and ADHD vulnerability (Cavenagh et al., 2019; Chatterjee et al., 2016), although activity levels in Xp22.31 deletion carriers have yet to be explicitly tested (e.g. using actigraphy). Future mouse work might examine the neurobiological basis of this hyperactivity phenotype, and, in particular, test whether it can be alleviated with medications used clinically to treat ADHD.

We found no significant main effects of genotype on any of the multiple anxiety-related behavioural measures we assessed. However, we did observe subtle, activity-independent, genotype by sex interactions for two such measures: open:closed arm time in the elevated plus maze, and willingness to consume a novel, and potentially toxic, solution. In both cases, homozygous male mice displayed behaviours consistent with lower anxiety (or greater risk-taking) than wildtype males (i.e. spending relatively more time on the open arms, and greater consumption of the novel foodstuff), whereas homozygous female mice displayed behaviours consistent with greater anxiety (or lower risk-taking) than wildtype females (i.e. spending relatively less time on the open arms, and lower consumption of the novel foodstuff). Although these findings could genuinely index effects on specific aspects of anxiety, it should be acknowledged that they could be Type I errors given our low power to reliably detect interactions, and the milk consumption data likely reflects hedonic (as well as anxiety-related or impulsivity-related) responses.

Our mouse findings contrast somewhat with findings in individuals with XLI, or female carriers, where elevated levels of generalised anxiety-related traits and anxiety disorder diagnoses are apparent (Wren et al., 2022). Perhaps the anxiety-related effects in man are due to loss of Xp22.31 genes other than *STS*, or perhaps STS has species-specific effects on anxiety processes. Testing individuals with XLI, or female carriers, in experimental paradigms more closely-related to those giving rise to significant findings in mice (e.g. virtual elevated plus maze (Biedermann et al., 2017)), might reveal analogous results.

For the remaining behavioural measures, indexing gross motor coordination (rotarod performance), reactivity to an acoustic stimulus and sensorimotor gating (startle response and inhibition of this upon exposure to a prepulse), and exploratory behaviour and working memory (spontaneous alternation), we observed no significant effects of genotype, or genotype interactions with sex. The fact that STS-deficient mice perform equivalently to wildtype mice in terms of these basic behaviours, means that they can be used subsequently without major issues for more sophisticated cognitive testing (e.g. assaying measure of attention and impulsivity associated with ADHD); however, the positive findings above (e.g. hyperactivity) will need to be accounted for in subsequent work.

Our endocrinological analysis identified the expected large sex differences in levels of testosterone and 5α-androstanedione, indicating that any large genotype-dependent effects on serum hormones should have been detectable. If genotype differences in serum hormone levels do genuinely exist, they are likely to be moderate, or small, in magnitude. In STS-deficient 39,X^Y*^O mice, we previously found no evidence for large changes in levels of brain steroids (Trent et al., 2014); however, these mice did exhibit significantly-lower levels of serum DHEA compared to 40,XY mice (Trent et al., 2012a). In the present analysis, median serum DHEA levels were lower, though not significantly so, in homozygous animals than in wildtype animals across both sexes.

Finally, we have shown that STS-deficient mice possess heavier hearts (normalised to bodyweight/tibia length) than their wildtype counterparts. Follow-up work will aim to understand the basis and sequelae of this result: structural analyses will test whether there is any evidence of predicted hypertrophy, while functional analyses will test for any associated effects on electrical signalling using electrocardiography (ECG). Analyses in the skin of STS-deficient mice have demonstrated altered Hippo signalling and elevated levels of phosphorylated YAP1 (Kwon et al., 2024). In cardiac cell culture, hyperactivation of YAP1 is associated with excessive CCN2 and TGFβ secretion and hypertrophy (Chirikian et al., 2024). Expression of these biochemical markers in STS-deficient mouse hearts may be perturbed, and warrants investigation. As cardiac screening in individuals with XLI becomes more commonly implemented, it will be interesting to see whether there is any evidence for hypertrophy in this group.

## 5. Conclusions

We have successfully generated a novel mouse model lacking STS activity, which exhibits normal breeding performance and is grossly-healthy. The model displays behavioural and cardiac phenotypes of relevance to XLI, but no large changes in serum steroid hormone biochemistry. Future work should aim to replicate the initial significant findings reported here, and to investigate the anatomical, cellular and molecular mechanisms underlying any robust results. More sophisticated phenotypic analyses, including of cognitive and cardiac function, might also be undertaken.

## Supporting information

Humby et al 2026 Supplemental Information

## Acknowledgements

Mice used in this study were obtained from the Mary Lyon Centre at MRC Harwell (MLC) and the following award is acknowledged: MC_UP_2201/2. The mouse model was generated in collaboration with Genome Editing Mice for Medicine Programme at Mary Lyon Centre MRC Harwell (https://www.har.mrc.ac.uk/projects/gemm/). Work was funded by an MRC GW4 DTP PhD studentship to Freya Shepherd, by a Cardiff University Innovation Development Scheme studentship to Talia Elgie, and by Cardiff University Schools of Psychology, Biosciences and Medicine. The funders played no role in the design, analysis or reporting of the study.

## References

Biedermann, S.V., Biedermann, D.G., Wenzlaff, F., Kurjak, T., Nouri, S., Auer, M.K., Wiedemann, K., Briken, P., Haaker, J., et al., 2017. An elevated plus-maze in mixed reality for studying human anxiety-related behavior. BMC Biol 15, 125. 10.1186/s12915-017-0463-6

Brcic, L., Underwood, J.F., Kendall, K.M., Caseras, X., Kirov, G., Davies, W., 2020. Medical and neurobehavioural phenotypes in carriers of X-linked ichthyosis-associated genetic deletions in the UK Biobank. J Med Genet 57, 692–98. 10.1136/jmedgenet-2019-106676

Bruter, A.V., Varlamova, E.A., Okulova, Y.D., Tatarskiy, V.V., Silaeva, Y.Y., Filatov, M.A., 2024. Genetically modified mice as a tool for the study of human diseases. Mol Biol Rep 51, 135. 10.1007/s11033-023-09066-0

Cavenagh, A., Chatterjee, S., Davies, W., 2019. Behavioural and psychiatric phenotypes in female carriers of genetic mutations associated with X-linked ichthyosis. PLoS One 14, e0212330. 10.1371/journal.pone.0212330

Chatterjee, S., Humby, T., Davies, W., 2016. Behavioural and Psychiatric Phenotypes in Men and Boys with X-Linked Ichthyosis: Evidence from a Worldwide Online Survey. PLoS One 11, e0164417. 10.1371/journal.pone.0164417

Chirikian, O., Faynus, M.A., Merk, M., Singh, Z., Muray, C., Pham, J., Chialastri, A., Vander Roest, A., Goldstein, A., et al., 2024. YAP dysregulation triggers hypertrophy by CCN2 secretion and TGFbeta uptake in human pluripotent stem cell-derived cardiomyocytes. bioRxiv. 10.1101/2024.06.03.597045

Davies, W., 2021. The contribution of Xp22.31 gene dosage to Turner and Klinefelter syndromes and sex-biased phenotypes. Eur J Med Genet 64, 104169. 10.1016/j.ejmg.2021.104169

Davies, W., 2025. The importance of cardiac screening in X-linked ichthyosis: a plea. Clin Exp Dermatol. 10.1093/ced/llaf221

Davies, W., Humby, T., Kong, W., Otter, T., Burgoyne, P.S., Wilkinson, L.S., 2009. Converging pharmacological and genetic evidence indicates a role for steroid sulfatase in attention. Biol Psychiatry 66, 360–7. 10.1016/j.biopsych.2009.01.001

Davies, W., Humby, T., Trent, S., Eddy, J.B., Ojarikre, O.A., Wilkinson, L.S., 2014. Genetic and pharmacological modulation of the steroid sulfatase axis improves response control; comparison with drugs used in ADHD. Neuropsychopharmacology 39, 2622–32. 10.1038/npp.2014.115

Fernandes, N.F., Janniger, C.K., Schwartz, R.A., 2010. X-linked ichthyosis: an oculocutaneous genodermatosis. J Am Acad Dermatol 62, 480–5. 10.1016/j.jaad.2009.04.028

Gilligan, L.C., Rahman, H.P., Tang, V., Hussain, M.T., Arvanti, A., Hewitt, A.M., Foster, P.A. 2017. Estrogen activation by steroid sulfatase increases colorectal cancer proliferation via GPER. J Clin Endocrinol Metab 102, 4435–4447. 10.1210/jc.2016-3716

GTEx Portal, 2025. https://www.gtexportal.org/home/gene/STS (accessed 20 June 2025)

Gurgen, D., Hegner, B., Kusch, A., Catar, R., Chaykovska, L., Hoff, U., Gross, V., Slowinski, T., da Costa Goncalves, A.C., et al., 2011. Estrogen receptor-beta signals left ventricular hypertrophy sex differences in normotensive deoxycorticosterone acetate-salt mice. Hypertension 57, 648–54. 10.1161/HYPERTENSIONAHA.110.166157

Kalev-Altman, R., Becker, G., Levy, T., Penn, S., Shpigel, N.Y., Monsonego-Ornan, E., Sela-Donenfeld, D., 2023. Mmp2 Deficiency Leads to Defective Parturition and High Dystocia Rates in Mice. Int J Mol Sci 24. 10.3390/ijms242316822

Kasahara, T., Mekada, K., Abe, K., Ashworth, A., Kato, T., 2022. Complete sequencing of the mouse pseudoautosomal region, the most rapidly evolving ‘chromosome’. bioRxiv 10.1101/2022.03.26.485930

Kutzler, M.A., 2008. Dystocia and Obstetric Crises, Small Animal Critical Care Medicine, 1st ed. Saunders, pp. 611–15. 10.1016/B978-1-4160-2591-7.10140-7

Kwon, T.U., Kwon, Y.J., Baek, H.S., Park, H., Lee, H., Chun, Y.J., 2024. Unraveling the molecular mechanisms of cell migration impairment and apoptosis associated with steroid sulfatase deficiency: Implications for X-linked ichthyosis. Biochim Biophys Acta Mol Basis Dis 1870, 167004. 10.1016/j.bbadis.2023.167004

Kwon, T.U., Kwon, Y.J., Park, H., Lee, H., Kwak, J.H., Kang, K.W., Chun, Y.J., 2025. Steroid sulfatase suppresses keratinization by inducing proteasomal degradation of E-cadherin via Hakai regulation. Biochim Biophys Acta Mol Cell Res 1872, 119898. 10.1016/j.bbamcr.2025.119898

Levenson, D., Romero, R., Garcia-Flores, V., Miller, D., Xu, Y., Sahi, A., Hassan, S.S., Gomez-Lopez, N., 2020. The effects of advanced maternal age on T-cell subsets at the maternal-fetal interface prior to term labor and in the offspring: a mouse study. Clin Exp Immunol 201, 58–75. 10.1111/cei.13437

Marx, J.O., Brice, A.K., Boston, R.C., Smith, A.L., 2013. Incidence rates of spontaneous disease in laboratory mice used at a large biomedical research institution. J Am Assoc Lab Anim Sci 52, 782–91.

Mianné, J., Codner, G.F., Caulder, A., Fell, R., Hutchison, M., King, R., Stewart, M.E., Wells, S., Teboul, L., 2017. Analysing the outcome of CRISPR-aided genome editing in embryos: Screening, genotyping and quality control. Methods 121-122, 68–76. 10.1016/j.ymeth.2017.03.016

Mouse Genome Informatics, 2025a. Genes and Markers. https://www.informatics.jax.org/marker/key/13669 (accessed 20 June 2025)

Mouse Genome Informatics, 2025b. Inbred strains of mice: C57BL. https://www.informatics.jax.org/inbred_strains/mouse/docs/C57BL.shtml (accessed 23 June 2025)

Mueller, J.W., Gilligan, L.C., Idkowiak, J., Arlt, W., Foster, P.A., 2015. The Regulation of Steroid Action by Sulfation and Desulfation. Endocr Rev 36, 526–63. 10.1210/er.2015-1036

Nicolas, L.B., Pinoteau, W., Papot, S., Routier, S., Guillaumet, G., Mortaud, S., 2001. Aggressive behavior induced by the steroid sulfatase inhibitor COUMATE and by DHEAS in CBA/H mice. Brain Res 922, 216–22. 10.1016/s0006-8993(01)03171-7

Perez-Jimenez, M.M., Monje-Moreno, J.M., Brokate-Llanos, A.M., Venegas-Caleron, M., Sanchez-Garcia, A., Sansigre, P., Valladares, A., Esteban-Garcia, S., Suarez-Pereira, I., et al., 2021. Steroid hormones sulfatase inactivation extends lifespan and ameliorates age-related diseases. Nat Commun 12, 49. 10.1038/s41467-020-20269-y

Rhodes, M.E., Li, P.K., Burke, A.M., Johnson, D.A., 1997. Enhanced plasma DHEAS, brain acetylcholine and memory mediated by steroid sulfatase inhibition. Brain Res 773, 28–32. 10.1016/s0006-8993(97)00867-6

Salido, E.C., Li, X.M., Yen, P.H., Martin, N., Mohandas, T.K., Shapiro, L.J., 1996. Cloning and expression of the mouse pseudoautosomal steroid sulphatase gene (Sts). Nat Genet 13, 83–6. 10.1038/ng0596-83

Schiffer, L., Shaheen, F., Gilligan, L.C., Storbeck, K-H., Hawley, J.M., Keevil, B.G., Arlt, W., Taylor, A.E., 2022. Multi-steroid profiling by UHPLC-MS/MS with post-column infusion of ammonium fluoride. J Chromatogr B Analyt Technol Biomed life Sci 1209, 123413. 10.1016/j.jchromb.2022.123413

The Jackson Laboratory, 2025. C57BL/6J. https://www.jax.org/strain/000664. (accessed 23 June 2025)

Trent, S., Fry, J.P., Ojarikre, O.A., Davies, W., 2014. Altered brain gene expression but not steroid biochemistry in a genetic mouse model of neurodevelopmental disorder. Mol Autism 5, 21. 10.1186/2040-2392-5-21

Trent, S., Dean, R., Veit, B., Cassano, T., Bedse, G., Ojarikre, O.A., Humby, T., Davies, W., 2013. Biological mechanisms associated with increased perseveration and hyperactivity in a genetic mouse model of neurodevelopmental disorder. Psychoneuroendocrinology 38, 1370–80. 10.1016/j.psyneuen.2012.12.002

Trent, S., Dennehy, A., Richardson, H., Ojarikre, O.A., Burgoyne, P.S., Humby, T., Davies, W., 2012a. Steroid sulfatase-deficient mice exhibit endophenotypes relevant to attention deficit hyperactivity disorder. Psychoneuroendocrinology 37, 221–9. 10.1016/j.psyneuen.2011.06.006

Trent, S., Cassano, T., Bedse, G., Ojarikre, O.A., Humby, T., Davies, W., 2012b. Altered serotonergic function may partially account for behavioral endophenotypes in steroid sulfatase-deficient mice. Neuropsychopharmacology 37, 1267–74.10.1038/npp.2011.314

Wren, G., Baker, E., Underwood, J., Humby, T., Thompson, A., Kirov, G., Escott-Price, V., Davies, W., 2023. Characterising heart rhythm abnormalities associated with Xp22.31 deletion. J Med Genet 60, 636–43. 10.1136/jmg-2022-108862

Wren, G.H., Davies, W., 2022. X-linked ichthyosis: New insights into a multi-system disorder. Skin Health Dis 2, e179. 10.1002/ski2.179

Wren, G.H., Davies, W., 2024. Cardiac arrhythmia in individuals with steroid sulfatase deficiency (X-linked ichthyosis): candidate anatomical and biochemical pathways. Essays Biochem 68, 423–29. 10.1042/EBC20230098

Wren, G.H., Flanagan, J., Underwood, J.F.G., Thompson, A.R., Humby, T., Davies, W., 2024. Memory, mood and associated neuroanatomy in individuals with steroid sulphatase deficiency (X-linked ichthyosis). Genes Brain Behav 23, e12893.10.1111/gbb.12893

Wren, G.H., Humby, T., Thompson, A.R., Davies, W., 2022. Mood symptoms, neurodevelopmental traits, and their contributory factors in X-linked ichthyosis, ichthyosis vulgaris and psoriasis. Clin Exp Dermatol 47, 1097–108. 10.1111/ced.15116

Zetoune, A.B., Fontaniere, S., Magnin, D., Anczukow, O., Buisson, M., Zhang, C.X., Mazoyer, S., 2008. Comparison of nonsense-mediated mRNA decay efficiency in various murine tissues. BMC Genet 9, 83. 10.1186/1471-2156-9-83

