## Supplementary material for "Physiological and behavioural characterisation of a novel steroid sulfatase-deficient mouse": Humby et al 2026 Supplemental Information

### Supplementary Information

#### Generation of *Sts*-deletion mice

Cas9 protein, single guide RNAs and single-stranded oligodeoxynucleotides (ssODNs) were diluted and mixed in Electroporation Buffer (Gibco Opti-MEM I Reduced Serum Media, Thermo Fisher Scientific) to working concentrations of 650 ng/µl, 130 ng/µl each and 400 ng/µl, respectively. Protospacer sequences were GGTGAAGCTGACGCAGCACC and GAAGCTGACGCAGCACCTGG with PAM sequences of TGG and CGG respectively. C57BL6/J one-cell stage embryos were electroporated using the following conditions: 30V, 3ms pulse length, 100ms pulse interval, 12 pulses. Electroporated embryos were implanted in CD1 pseudo-pregnant females. Host females were allowed to litter and rear $F_0$ progeny. Genomic DNA was extracted from ear biopsies of progeny and the presence of the anticipated genetic change (a 22 basepair sequence (TGACGCAGCACCTGGCGGCCGC) deletion from the sulfatase domain-encoding exon 2, resulting in a frameshift mutation) was tested for using the following PCR conditions: F:GAGGCCTGACACAGGCAG, R: GTAGCGCCCGGTCAGGAA, annealing temperature 60°C and an elongation time of 15s with amplicons sent for Sanger sequencing. One off-target exonic site (2:119,169,717-119,169,739) with ≤2 mismatches for guide(s) used was checked with the following primers: F:GCTTCAGTTTCACGAACGGT and R: TCCTAACGCTCAAGAGTGGG and amplicons sent for Sanger sequencing. No off-target activity was detected in animals used to establish the colony.

#### Enzyme activity assay methods

Frozen tissue (100 mg wet weight) was combined with liquid nitrogen and broken up using a pestle and mortar. The resultant homogenate was then stored at -80°C. Approximately 2-5mg of tissue sample was then added to a glass tube. Enzyme reactions (500µl final volume) containing 0.1% BSA and up to 1µM [6,7-$^{3}$H]oestrone sulfate (≈1×10$^{6}$ dpm) were prepared. Tissue protein was added to these, and tubes were incubated at 37°C for 18hrs with occasional gentle shaking. Reactions were quenched on ice with 4ml toluene, vortex-mixed for 30s and centrifuged at 3000 g for 10 mins. The organic phase was then removed for analysis. STS activity (nmol/mg/h) was calculated from the amount of oestrone formed, normalised to wet weight and to incubation time.

#### RNA extraction and *Sts* gene expression analysis methods

Liver samples (~10mg) were lysed in 1:100 2-mercaptoethanol in RLT buffer using stainless steel lysis beads (Qiagen, 69989) in a TissueLyser II (Qiagen). RNA was subsequently extracted using the Qiagen RNeasy Mini Kit (74104) as per manufacturer's instructions and eluted into RNase-free water. RNA quality and quantity was assessed using a NanoDrop spectrophotometer (ThermoFisher Scientific). cDNA was generated from high-quality RNA using High-Capacity cDNA Reverse Transcription Kit (Applied Biosystems, 4368814), and all samples were normalised to 100ng/ml. Gene expression was determined using the following TaqMan Gene Expression Assay probes (ThermoFisher Scientific): *Gapdh* (Mm99999915_g1) and *Sts* (Mm04214605_u1) with a QuantStudio 5 Real-Time PCR system (Applied Biosystems, Warrington, UK). *Sts* expression was normalised to *Gapdh* expression (ΔCt) and the fold-difference presented using arbitrary units (AU = 1000 x ($2^{-\Delta Ct}$)). Expression levels across genotypes were compared by unpaired t-test (**p<0.01)

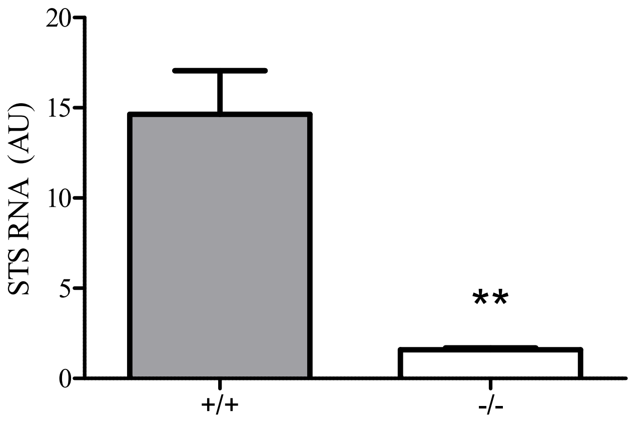


**Supplementary Figure 1.** *Sts* gene expression patterns in liver tissue of wildtype (+/+) and homozygous (-/-) male and female mice

### Steroid hormone assay methods

Steroids were extracted from 1-200µL of serum after addition of 20µL of an internal standard mixture. Steroids were extracted via liquid/liquid extraction with MTBE (tert-methyl butyl ether). The organic layer was removed into a 96-well plate where it was evaporated under a stream of heated nitrogen (50°C). Extracts were reconstituted in 125µl of 50:50 methanol:water for analysis using liquid chromatography-mass spectrometry. Steroid quantification was achieved through comparison to a calibration series on a Waters Xevo-XS triple quadruple mass spectrometer with Acquity uPLC.

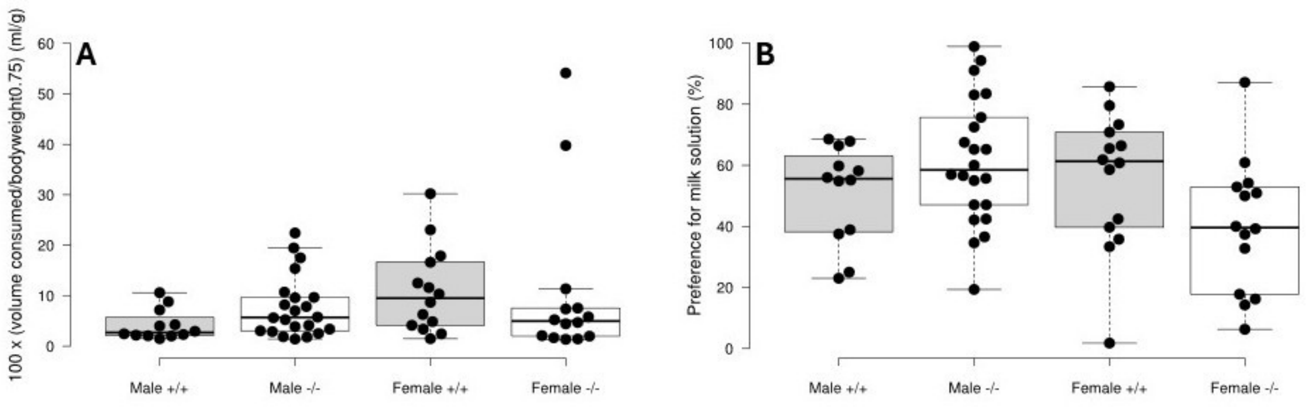


**Supplementary Figure 2.** Performance of adult male and female wildtype (+/+) and homozygous (-/-) mice on key measures of consummatory behaviour (volume consumed per bodyweight (A) and preference for milk over water (B)) on the milk preference test

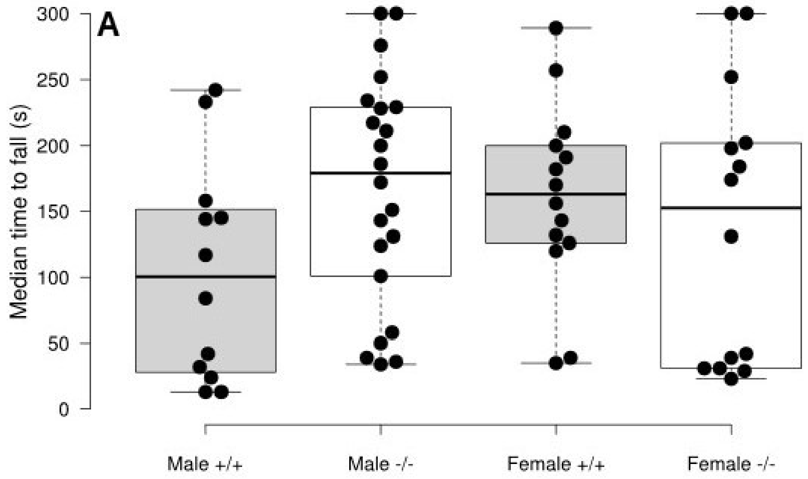


**Supplementary Figure 3.** Performance of adult male and female wildtype (+/+) and homozygous (-/-) mice on the rotarod test (median time to fall from an increasingly rapidly rotating rod across 5 individual trials)

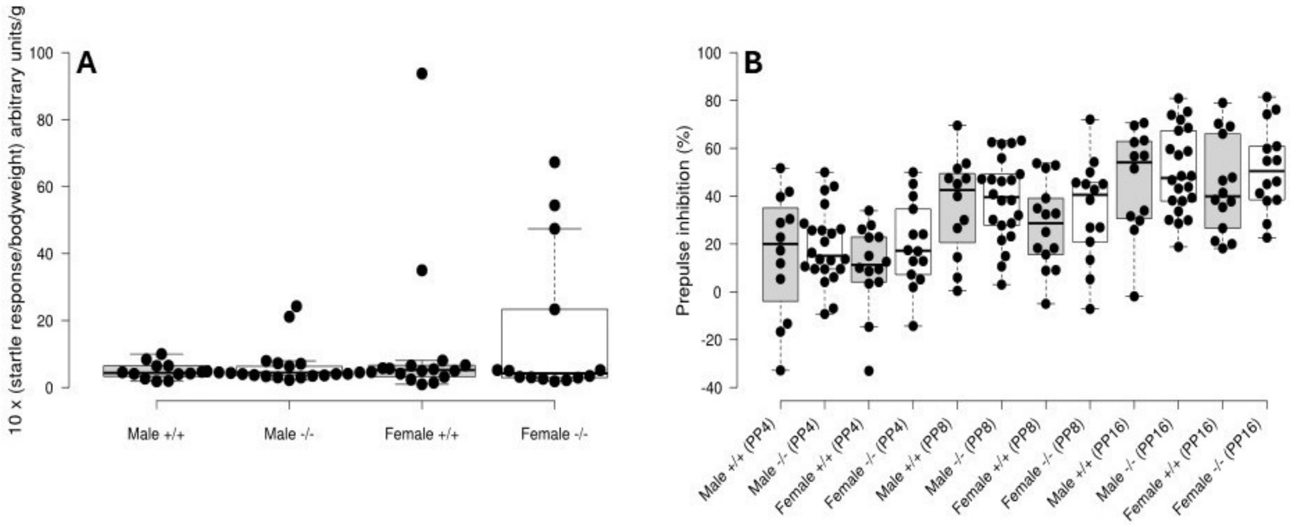


**Supplementary Figure 4.** Performance of adult male and female wildtype (+/+) and homozygous (-/-) mice on the startle (A) and prepulse inhibition (B) test. Prepulses of 4, 8 and 16dB were used (PP4, PP8 and PP16 respectively).
